# Dispersal ability modulates macroinvertebrate metacommunity responses to environmental filtering and spatial connectivity in an Afrotropical stream network

**DOI:** 10.64898/2026.07.30.741733

**Authors:** Bashizi C. Christian, Elizabeth W. Wanderi, Augustine Sitati, Mourine J. Yegon, Thomas Fuß, Frank O. Masese

## Abstract

**Aim:** To test whether dispersal ability determines the relative importance of environmental filtering and distinct dispersal pathways in structuring stream macroinvertebrate metacommunities.

**Location:** Nzoia River catchment, Lake Victoria basin, western Kenya.

**Taxon:** Benthic macroinvertebrates (family level), grouped by dispersal ability into strong fliers, weak fliers, passive aerial dispersers, and obligate aquatics.

**Methods:** Macroinvertebrates were sampled seasonally (2018 to 2021) at 55 sites. Using distance-based redundancy analysis and variation partitioning, we compared the influence of physicochemical water quality on community structure, representing environmental filtering, with the influence of three sets of spatial factors representing distinct dispersal pathways: Euclidean distance (overland dispersal pathway), land-use resistance (land-use mediated dispersal pathway), and Asymmetric Eigenvector Maps built on river-network connectivity (along-watercourse dispersal pathway). In addition, distance-decay relationships and turnover-nestedness partitioning were examined for the full assemblage and each dispersal group.

**Results:** Community assembly shifted predictably along the macroinvertebrate dispersal-ability gradient. Environmental filtering dominated in strong fliers and retained an independent influence in weak fliers and obligate aquatics, but contributed no independent influence in passive aerial dispersers. The overland dispersal pathway structured the three aerial groups. The land use mediated dispersal pathway constrained weak fliers and passive aerial dispersers, and the along-watercourse dispersal pathway structured passive aerial dispersers and obligate aquatics. Passive aerial dispersers showed the strongest dispersal signal, with the overland and along-watercourse pathways contributing equally, and in this group beta diversity reflected richness gradients (nestedness) rather than species replacement.

**Main conclusions:** Dispersal ability determined whether environmental filtering or a specific dispersal pathway governed community composition, and no single pathway captured assembly across the full range of dispersal abilities. Dispersal traits and connectivity must therefore be considered jointly when studying and managing aquatic biodiversity in human-modified Afrotropical catchments.

## Introduction

Stream macroinvertebrates are central to freshwater biodiversity and ecosystem functioning, but many taxa also depend on terrestrial corridors for adult dispersal (Wallace & Webster, 1996; Bilton et al., 2001; Hauer & Resh, 2017). Movement among habitat patches supports recolonization, population connectivity, and gene flow, thereby maintaining the persistence and diversity of populations across stream networks (Paetzold et al., 2005; Lopes et al., 2025; Burdon et al., 2025). Their community composition therefore reflects both local instream conditions that filter larvae and the spatial permeability of the landscape through which adults disperse (Carlson et al., 2016; Cano-Barbacil et al., 2025). In catchments modified by agriculture, urbanization and riparian degradation, determining the relative influence of local conditions and landscape permeability is essential because water-quality restoration and connectivity restoration target different parts of the life cycle.

Land use change alters streams through nutrient enrichment, sedimentation, channel simplification, loss of canopy cover and altered thermal regimes, with well-documented consequences for macroinvertebrate taxonomic and functional composition (Naiman et al., 2010; Masese et al., 2014; Chen et al., 2023; Rumschlag et al., 2023). Local in-stream conditions regulate larval survival, their development, and emergence, whereas the terrestrial adult phase governs dispersal, reproduction and connectivity while exposing adults to habitat fragmentation, altered microclimates, loss of riparian vegetation and disconnected migration corridors, and barriers to movement (Bunn & Hughes, 1997; Leipold et al., 2004; Roodt et al., 2023). Macroinvertebrate community structure therefore reflects not only local habitat quality within stream reaches but also the degree to which the landscape facilitates or constrains movement among sites (Firmiano et al., 2021; Li et al., 2021). In human-modified catchments where landuse heterogeneity is pronounced and riparian buffers are often degraded, a metacommunity analysis that incorporates alternative connectivity models could therefore be stronger than one that treats spatial structure as a single undifferentiated distance effect.

Metacommunity theory links local environmental filtering with regional dispersal (Leibold et al., 2004; Cottenie, 2005). Environmental filtering drives species sorting, where taxa occur at sites with suitable environmental and habitat conditions. Community assembly processes include dispersal limitation, where restricted movement generates spatial structure independent of the measured environment, and mass effects, where high immigration sustains taxa outside their environmental optima (Heino et al., 2015; Tonkin et al., 2018). We therefore distinguish assembly processes (environmental filtering versus dispersal limitation or mass effects) from dispersal pathways, the physical routes through which organisms move. Beta diversity helps interpret these processes because turnover indicates compositional replacement along environmental or spatial gradients, whereas nestedness indicates richness gradients and potential homogenizing effects of dispersal (Whittaker, 1962; Baselga, 2010).

Straight-line (Euclidean) geographic distance poorly represents the effective separation between sites, because macroinvertebrates migrate through landscapes that vary in land use composition and river-network configuration (Smith et al., 2015; Morán-Ordóñez et al., 2015; Sarremejane et al., 2017; de Araujo Barbosa et al., 2025). Land use resistance captures the permeability of the terrestrial matrix to adult dispersal, whereas the dendritic structure of river networks confines aquatic dispersers (mainly larvae) to the river channel, so both provide more realistic connectivity measures than Euclidean distance (Petersen et al., 2004; Firmiano et al., 2021). Combining environmental and spatial predictors has improved predictions of community composition and assembly in temperate and Neotropical systems (Morán-Ordóñez et al., 2015; Armendáriz et al., 2022; Rodríguez et al., 2022). However, the relevant spatial predictor depends on dispersal ability: obligate aquatic taxa that lack an aerial adult stage are structured mainly by river-network connectivity, whereas aerial dispersers are structured by land-use resistance reflecting matrix permeability and geographic distance between sites (Liu et al., 2013; Tonkin et al., 2018; Richmond et al., 2024). Pooling all taxa therefore obscures how environmental filtering and spatial processes jointly shape assemblages.

Across Afrotropical stream networks, agricultural expansion and intensification, deforestation, urbanization, and poor wastewater management increasingly alter water quality, habitat structure and the structural and functional organization of stream communities (Masese et al., 2017; Fugère et al., 2018; Yegon et al., 2021; Akamagwuna et al., 2022; Dalu & Masese, 2024). The Nzoia River catchment offers a suitable case study, draining approximately 12,900 km² of western Kenya into Lake Victoria across forested, agricultural, mixed and urban land uses, and degraded riparian zones (Khan et al., 2011; Sitati et al., 2021). Here, local physicochemical conditions should strengthen environmental filtering by favouring most adapted taxa, whereas reduced connectivity should increase dispersal limitation, particularly among weak dispersers; species-sorting should be most detectable where connectivity permits colonization but is not so high that mass effects override local selection (Leibold et al., 2004; Lindström & Langenheder, 2012; Heino et al., 2015; Tonkin et al., 2018). Although environmental filtering and dispersal are recognized joint drivers of stream metacommunities, evidence remains scarce for Afrotropical networks, particularly on whether Euclidean distance, land-use resistance and river-network connectivity differ in explanatory power across dispersal groups.

To address this gap, we examined how different dispersal pathways shape assembly processes in an Afrotropical riverine macroinvertebrate metacommunity. We evaluated how environmental predictors, comprising physicochemical water quality and river longitudinal gradients (depth, velocity, wetted width), and three spatial predictors structure macroinvertebrate community composition across the Nzoia River catchment. Each spatial predictor represents a distinct dispersal pathway: Euclidean distance represents the overland dispersal pathway, land-use resistance the land-use mediated dispersal pathway, and Asymmetric Eigenvector Maps built on river-network connectivity the along-watercourse dispersal pathway. The specific objectives were: (i) to quantify the relative importance of environmental and spatial predictors on macroinvertebrate community composition; (ii) to compare the explanatory power of the three dispersal pathways for community composition; (iii) to test whether the explanatory power of environmental and spatial predictors varies among dispersal groups; and (iv) to determine whether community dissimilarity increases along environmental and spatial distance gradients within the full assemblage and within each dispersal group. We hypothesised (H1) that assemblage structure is jointly shaped by local environmental filtering and by regional dispersal, which determines which taxa migrate across the catchment. We further hypothesised (H2) that the relative importance of environmental filtering and dispersal pathways differs among dispersal groups: strong fliers are shaped primarily by local environmental filtering, with minimal independent contribution from land use resistance; weak fliers are more constrained by the land use resistance than strong fliers, because their short female dispersal distances make the terrestrial matrix a relevant barrier to successful dispersal; passive aerial dispersers are shaped primarily by dispersal downstream along the watercourse and potentially, airflow along the stream valley corridor (Vanschoenwinkel et al., 2009); and obligate aquatics are structured by both dispersal downstream along the watercourse and local environmental filtering, because instream physicochemical conditions provide the only habitat filter for taxa with no aerial adult stage. We also hypothesised (H3) that community dissimilarity increases with increasing environmental and effective spatial distance, with stronger distance-decay relationships among dispersal-limited groups than among strong fliers, and with the steepest along-watercourse decay for obligate aquatics.

## Materials and Methods

### Study area

The Nzoia River basin (Figure 1) lies in western Kenya, fed by streams draining three major water towers: Mt Elgon, the Cherangani Hills, and the Mau Forest. The basin drains an area of about 12,900 km² and discharges into Lake Victoria (Odira et al., 2010; Maloba et al., 2016; Rotich et al., 2022). Topography, climate, hydrology, and land use vary along its length, from forested headwaters through agricultural middle reaches to mixed lower plains. The climate is mainly tropical humid, with mean annual rainfall ranging from 900 to 2,200 mm and mean monthly temperature ranging from 13 to 25 °C, both varying strongly with elevation and season (Kadeka et al., 2021). Rainfall is highest in the headwaters and decreases towards the lowlands (Musau et al., 2015). The annual rainfall pattern is bimodal, with long rains from March to June and short rains from August to October (Kadeka et al., 2021). The Nzoia River flows throughout the year but shows a strong seasonal increase in discharge during the two rainy seasons, often flooding its floodplains (Immerzeel & Droogers, 2009; Akali et al., 2015; Moses et al., 2015). Land use change through deforestation, cultivation, livestock grazing, and the expansion of settlements and urban areas have altered the natural flow regime, erosion, and sediment delivery, producing a heterogeneous landscape suited to testing how local water quality and landscape connectivity jointly structure stream communities (Githui et al., 2009; Maloba et al., 2016; Koskei & Janeth, 2024). Grassland (45.82%) and cropland (38.52%) dominate land cover, followed by forest (14.13%). Settlements and urban areas (1.22%), bareland (0.22%), and water bodies (0.08%) occupy small fractions (Figure 1).

**Figure 1.**
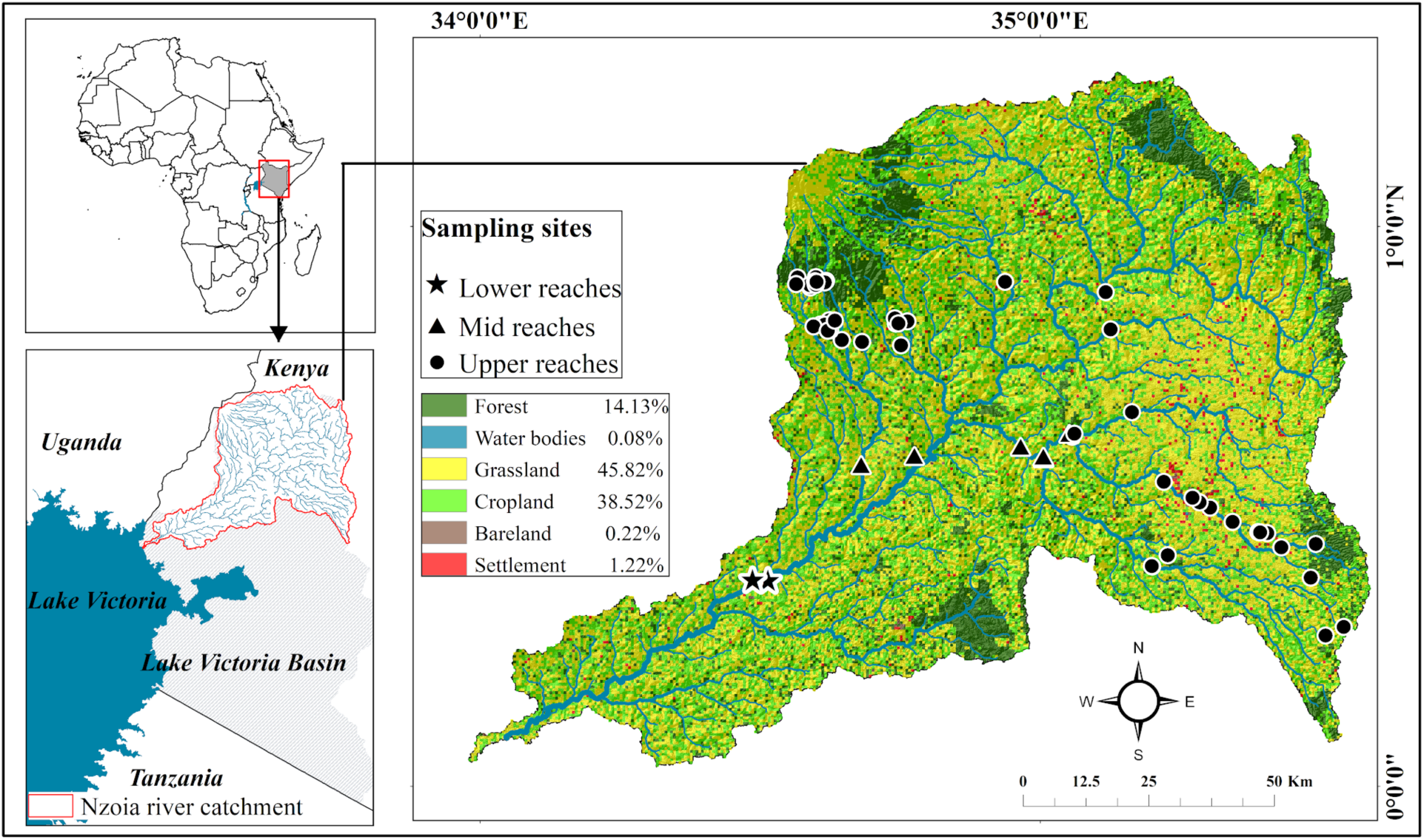
Location of the Nzoia River catchment in western Kenya, showing the distribution of the 55 macroinvertebrate sampling sites and the spatial pattern of major land use classes. Insets show the geographical position of the catchment within the Lake Victoria Basin and Africa.

**Figure 2.**
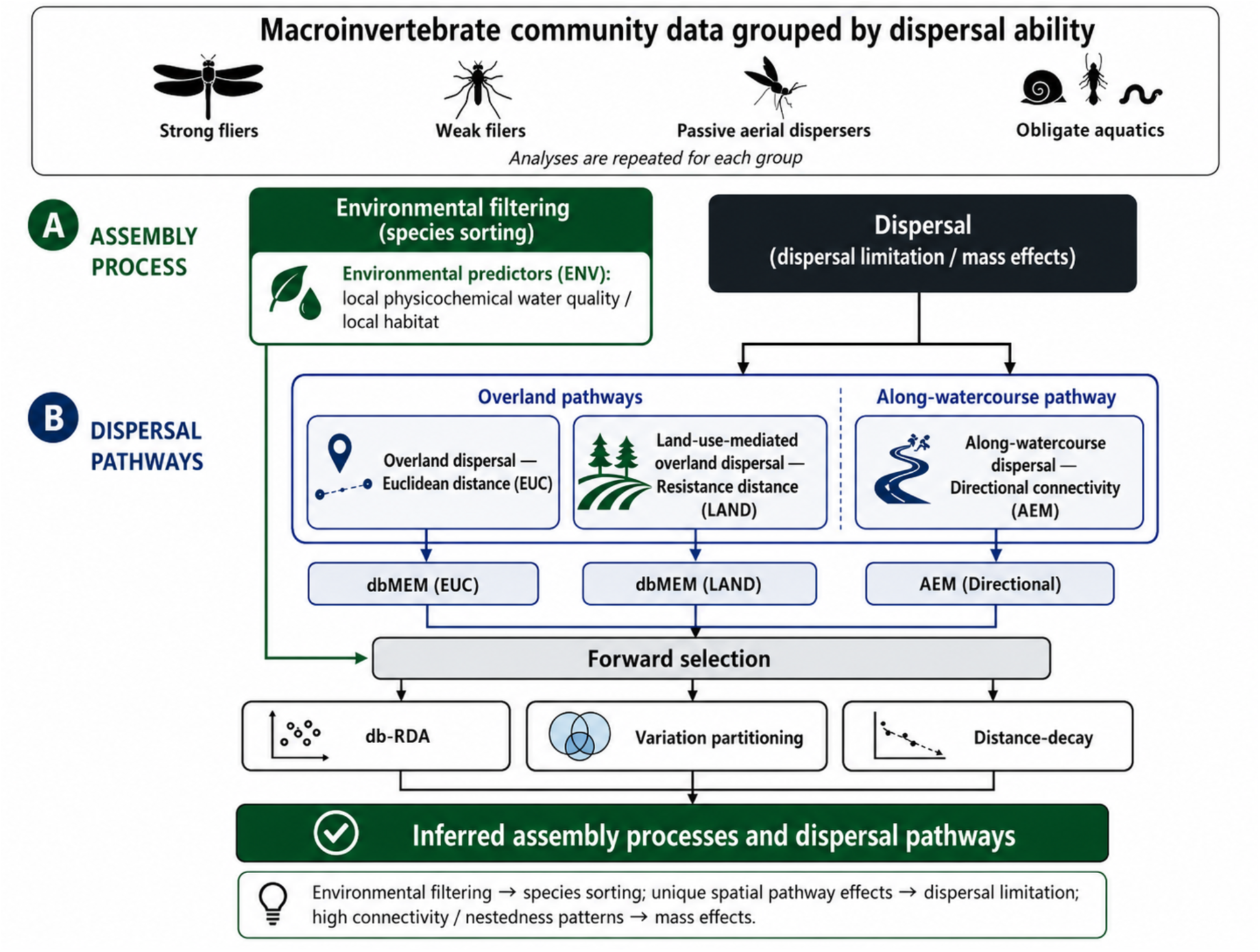
Conceptual framework separating assembly processes from dispersal pathways for macroinvertebrate metacommunities. Environmental predictors (ENV) represent environmental filtering, which drives species sorting, whereas the spatial factors Euclidean distance (EUC), land-use resistance (LAND) and Asymmetric Eigenvector Maps built on river-network connectivity (AEM) represent the overland, land-use mediated and along-watercourse dispersal pathways, respectively. Analyses were repeated for each dispersal group using db-RDA, variation partitioning and distance-decay models.

**Figure 3.**
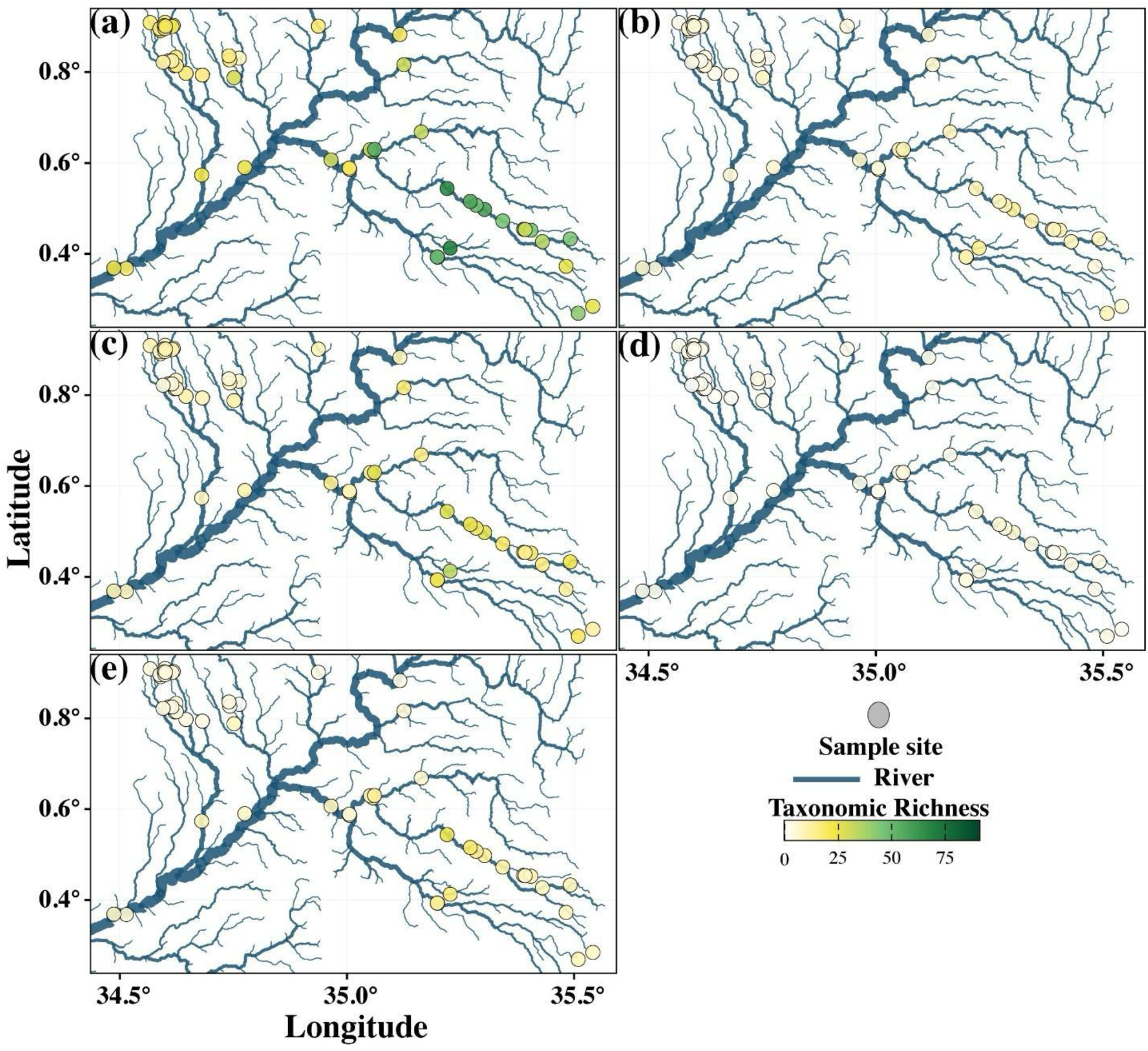
Spatial distribution of macroinvertebrate taxonomic richness across the 55 sampling sites. Richness is shown for the full assemblage (a) and the four dispersal groups: (b) strong fliers, (c) weak fliers, (d) passive aerial dispersers, (e) obligate aquatics. Symbol colour scales with family richness per site.

**Figure 4.**
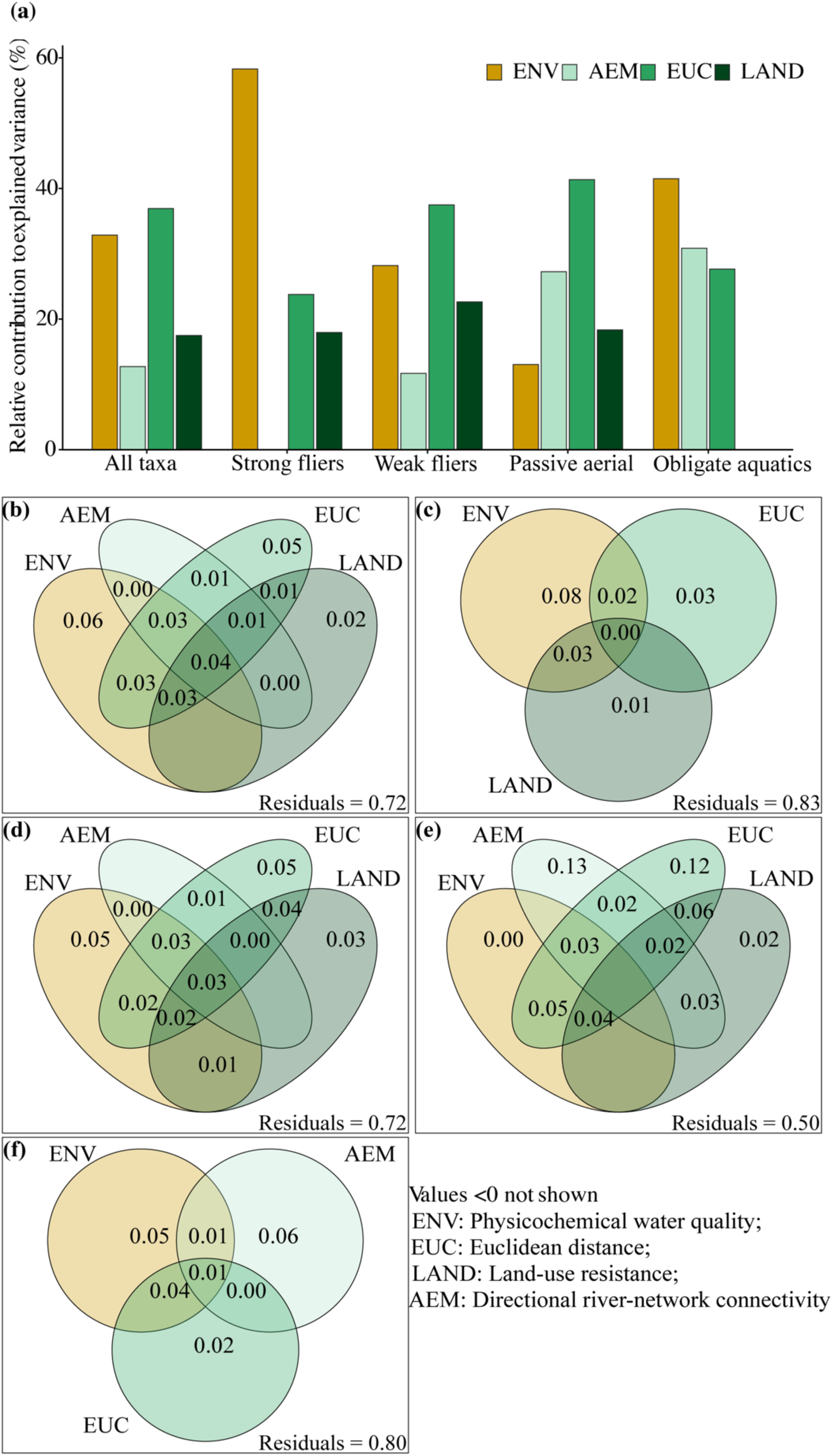
Predictor block contributions to macroinvertebrate community composition for the full assemblage and four dispersal groups. Panel (a) shows each block’s adjusted R² as a percentage of the summed adjusted R² of blocks significant at *p* ≤ 0.05. Panels (b) to (f) show variation partitioning as adjusted R² fractions for the full assemblage (b), strong fliers (c), weak fliers (d), passive aerial dispersers (e), and obligate aquatics (f). Shared fractions reflect correlation among blocks and are not attributable to any single block. Residuals denote unexplained variance.

### Macroinvertebrate sampling and identification

Macroinvertebrates were sampled seasonally from 2018 to 2021 at 55 stream sites using a semi-quantitative kick-net procedure based on SASS5 protocol (Dickens & Graham, 2002). At each 100 m reach, the main biotopes (gravel, sand/mud, stones in current and stones out of current, marginal vegetation and litter packs/ coarse organic matter) were sampled with four replicate 500 μm kick-net samples per biotope. Within each reach, sampling proceeded from downstream to upstream to minimize disturbance and drift. Samples were field-sorted, preserved in 70% ethyl ethanol and transported to the laboratory. Samples were rinsed through a 500 µm sieve to remove fine debris, transferred to sorting trays. All individuals were enumerated and identified to genus level whenever possible using standard taxonomic keys and guides, including Fry (2022), Day & De Moor (2002), de Moor & Day (2003), and Merritt et al. (2008). For the present study, however, all analyses were conducted at family level.

### Classification of dispersal groups

Macroinvertebrate families were classified into dispersal groups following Morán-Ordóñez *et al*. (2015) and Firmiano *et al*. (2021). Classification was based on two functional traits, namely adult flying strength and female dispersal distance, yielding four groups: obligate aquatics, passive aerial dispersers, weak fliers, and strong fliers (Table S1). Although species-level classification would be preferable, dispersal capacity tends to be conserved within families (Poff et al., 2006), and limited species-level trait information was available for the Afrotropical region, justifying a family-level approach. Based on this classification, we analyzed the full assemblage and each dispersal group separately.

### Measurements of physicochemical variables

At each site, before sampling macroinvertebrates, *in-situ* measurements of water quality parameters such as dissolved oxygen (DO; mg L⁻¹), temperature (°C), electrical conductivity (EC; μS cm⁻¹), salinity (ppt) and pH were recorded using a YSI multiprobe water quality meter (556 MPS; Yellow Springs Instruments, Yellow Springs, OH, USA). Stream width was measured at 10 equally spaced transects within 100 m of a sampling reach. At each transect, water depth and flow velocity were determined using a velocity plank, and stream discharge was calculated using the velocity–area method (Wetzel & Likens, 2000). To measure total suspended solids (TSS) and particulate organic matter (POM), triplicate water samples of known volumes of water were filtered through pre-combusted and pre-weighed Whatman GF/F glass fiber filters (pore size 0.7 μm, diameter 47 mm). The filters were wrapped in aluminum foil, stored in a cooler, and transported to the laboratory for further processing.

### Environmental and spatial predictors

To represent the main processes expected to structure community composition, four distance metrics were considered. One metric described local environmental conditions in terms of environmental distance and represented environmental filtering. The remaining three described spatial separation among sites and represented distinct dispersal pathways, namely Euclidean distance (EUC), land-use resistance (LAND), and Asymmetric Eigenvector Maps built on river-network connectivity (AEM) (Morán-Ordóñez et al., 2015; Sarremejane et al., 2017; Firmiano et al., 2021).

### Environmental conditions

Environmental distance represented differences in local physicochemical conditions among sites, serving as the environmental predictor (ENV) block in the analytical framework. Physicochemical variables were screened before analysis by removing near-zero-variance variables, one variable from each pair with |r| ≥ 0.7, and variables with VIF > 10, and then standardising the retained variables to zero mean and unit variance (Supporting Information, Appendix S1.1). Forward selection with dual stopping criteria, a permutation-based significance threshold of α = 0.05 and an adjusted R² ceiling equal to that of the full model, was then applied independently for each dispersal group to identify the subset of variables that significantly and non-redundantly explained variation in community composition (Blanchet et al., 2008a).

### Spatial predictors

Euclidean distance represented the straight-line geographic distance between sampling sites, calculated from their coordinates (Jones et al., 2010; Borcard et al., 2018). This distance matrix served as the input for the distance-based Moran Eigenvector Map (dbMEM) procedure (Legendre & Legendre, 2012), which we used to derive spatial predictors reflecting the hypothesis that dispersal declines with increasing geographic separation (Borcard et al., 2018). The resulting dbMEM eigenvectors in the Euclidean distance block (EUC) were derived exclusively from site coordinates and therefore represent undifferentiated overland spatial structure, independent of land use composition (Supporting Information Figure S2b).

Land use resistance was included as a separate spatial metric because variation in land use influences landscape permeability and therefore the effective dispersal distance among stream sites (Kärnä et al., 2015; Sarremejane et al., 2017; Firmiano et al., 2021; de Araujo Barbosa et al., 2025). The resistance surface was derived from Sentinel-2 imagery (10 m resolution; ESA, 2015) processed in Google Earth Engine (Gorelick et al., 2017), using a median composite of all available imagery acquired between 2018 and 2021 to produce a temporally representative characterization of land use across the sampling period. Raster cells were then assigned resistance values according to their expected effect on the movement of flying aquatic insects, with raster preparation, reclassification, and formatting conducted in ArcMap prior to analysis in CIRCUITSCAPE 4.0 (McRae et al., 2009). Resistance values were assigned to the dominant land use classes in the study area, namely forest, grassland, cropland, and urban areas, following Richmond et al. (2024) and expert-based ranking informed by published evidence on adult aquatic-insect dispersal (Smith et al., 2015; Carlson et al., 2016).

Asymmetric Eigenvector Maps (AEM) represent river-network connectivity and encode the asymmetry imposed by flow, in which connectivity runs mainly from upstream to downstream sites (Blanchet et al., 2008a, 2008b; Landeiro et al., 2011). We derived the river network from the 30 m SRTM DEM (Farr et al., 2007), providing the topographic basis for stream mapping and directional connectivity among sites (Supporting Information Figure S2c). Spatial eigenvectors for all three spatial factors were filtered to retain only those with significant positive spatial autocorrelation, computed with the moran.I.multi function in the *adespatial* package in R (Dray et al., 2006). We computed all spatial eigenvectors, dbMEM and AEM, in R with the adespatial package. Construction of the resistance surface and river-network eigenvectors is detailed in Supporting Information (Appendix S1.2 and S1.3).

## Statistical analysis

Community composition was analysed with distance-based redundancy analysis (dbRDA) using Bray-Curtis dissimilarities for the full assemblage and each dispersal group (Legendre & Anderson, 1999). Candidate ENV, EUC, LAND and AEM predictors were forward-selected by the same procedure, applied independently to each block and to each dispersal group. Selection used 999 permutations with dual stopping criteria, a permutation-based significance threshold of α = 0.05 and an adjusted R² ceiling equal to that of the full block model, to reduce overfitting (Blanchet et al., 2008a; Peres-Neto & Legendre, 2010). Analyses were implemented in R using vegan and *adespatial* packages.

Variation partitioning was then used to quantify the unique and shared contributions of the four predictor blocks (ENV, EUC, AEM, LAND) to changes in community composition (Borcard et al., 1992; Peres-Neto et al., 2006; Dray, 2012). The unique fraction of each block represents variance explained by that block after statistically controlling for all other blocks. The shared fractions reflect collinearity among predictors, and cannot be attributed to any single predictor independently (Diniz-Filho et al., 2012; Mouton et al., 2022). Variation partitioning was performed separately for all dispersal groups using functions available in R package *vegan*.

We analysed distance-decay relationships (Nekola & White, 1999) to test how community dissimilarity changed with environmental and spatial separation among sites (Morán-Ordóñez et al., 2015). We related pairwise Bray-Curtis dissimilarity to four distance metrics: environmental distance (Euclidean distance across standardised physicochemical variables; Legendre & Anderson, 1999; Peres-Neto & Legendre, 2010), geographic distance (Euclidean), resistance distance (CIRCUITSCAPE), and directional river network connectivity (AEM). Mantel tests (999 permutations) tested each relationship first. Linear regressions then quantified the direction and strength of each distance-dissimilarity relationship for the full assemblage and for each dispersal group.

Beta diversity was decomposed into turnover and nestedness components using functions in the R package *betapart* (Baselga, 2010; Baselga & Orme, 2012). Family-level abundances were converted to presence-absence, and total Sorensen dissimilarity was partitioned with beta.multi and beta.pair into a turnover component, the Simpson dissimilarity (beta.SIM) reflecting compositional replacement among sites, and a nestedness component (beta.SNE) reflecting richness differences among sites. The multi-site partition produced a single summary value per dispersal group, from which turnover and nestedness were expressed as percentages of total Sorensen dissimilarity (beta.SIM/beta.SOR and beta.SNE/beta.SOR). The pairwise partition gave the distributions of dissimilarity among site pairs. The decomposition was applied to the full assemblage and to each dispersal group separately.

## Results

### Macroinvertebrates composition

All taxa combined yielded the highest total abundance (1,036.2 ± 1,013.3 individuals per sample site, mean ± SD) and richness (22.5 ± 6.3 families per sample site) (Figure S1). Among individual dispersal groups, weak fliers dominated in both mean abundance (618.4 ± 542.3) and richness (10.2 ± 3.1), indicating that this group is the most numerically abundant and taxonomically diverse component of these communities. Strong fliers and passive aerial dispersers showed comparable mean abundances (104.4 ± 140.1 and 105.5 ± 280.4, respectively) but differed in richness, with strong fliers supporting more families per site (4.8 ± 1.9) than passive aerial taxa (1.9 ± 0.7). Obligate aquatics exhibited intermediate abundance (208.0 ± 637.9) and richness (5.5 ± 2.2).

### Influence of spatial and environmental factors

Three of the four predictor blocks explained significant proportions of community variance in most dispersal groups (Table 3). In the total dbRDA models, ENV was significant for every group (R²adj = 0.101 to 0.186, *p* ≤ 0.001), with the strongest explanatory power recorded for all taxa combined (R²adj = 0.186) and the weakest for passive aerial dispersers (R²adj = 0.101). Euclidean distance (EUC) was the dominant spatial factor for all taxa (R²adj = 0.209), weak fliers (R²adj = 0.202), and passive aerial dispersers (R²adj = 0.320). Asymmetric Eigenvector Maps built on river-network connectivity (AEM) were not selected for strong fliers, reached their highest value across groups in passive aerial dispersers (R²adj = 0.211) and explained a smaller proportion among obligate aquatics (R²adj = 0.065). Land-use resistance (LAND) explained significant variance in all groups except obligate aquatics (R²adj = 0.040 to 0.142, *p* ≤ 0.020), with the largest effect recorded for passive aerial dispersers (R²adj = 0.142) and weak fliers (R²adj = 0.122).

**Table 3.**
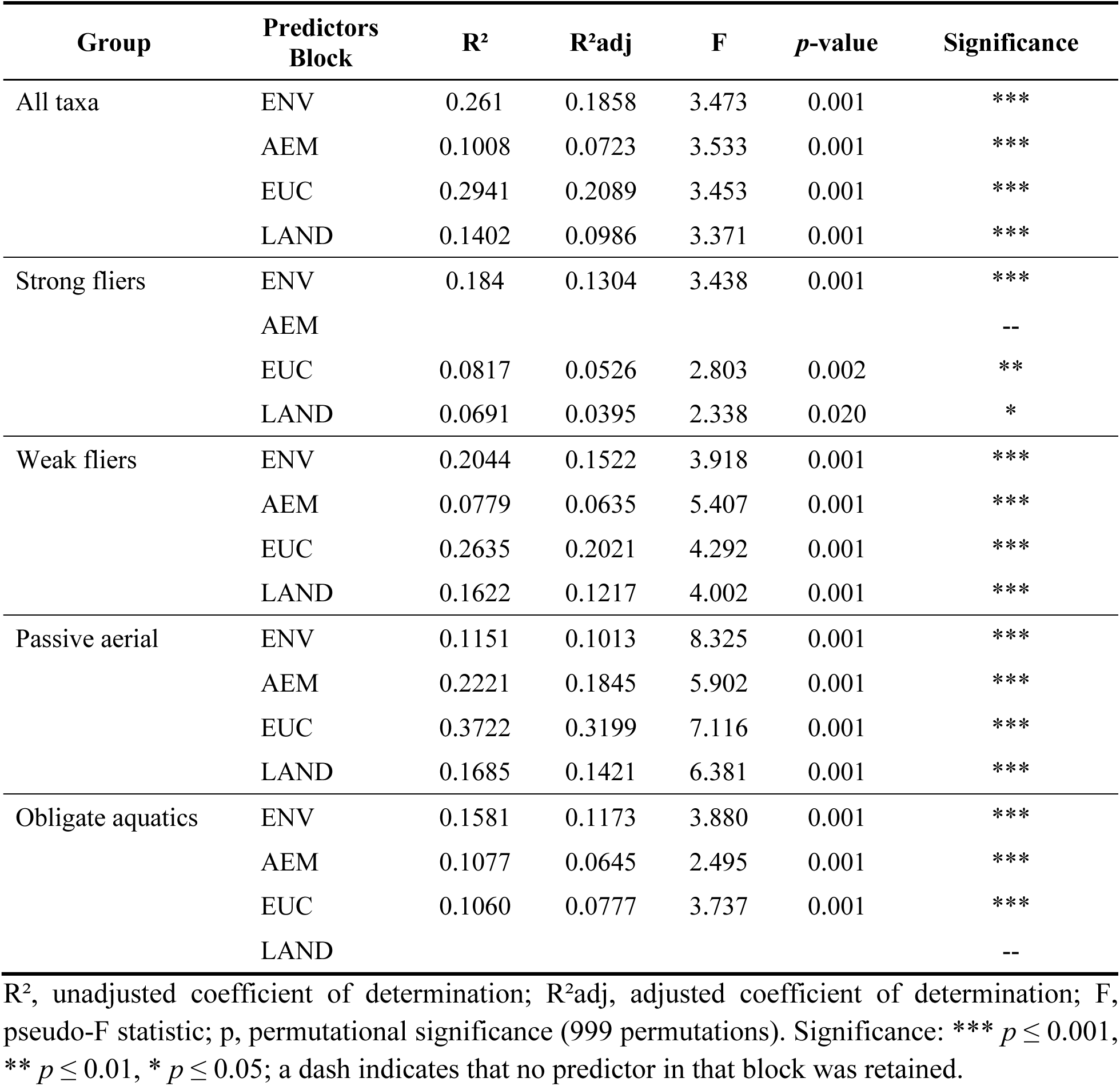
Distance-based redundancy analysis (db-RDA) of macroinvertebrate community composition against each predictor block. Models were fitted separately for the full assemblage and the four dispersal groups. ENV, physicochemical water quality; EUC, Euclidean distance; LAND, land-use resistance; AEM, river-network connectivity.

| Group | Predictors Block | R<sup>2</sup> | R<sup>2</sup><sub>adj</sub> | F | p-value | Significance |
| --- | --- | --- | --- | --- | --- | --- |
| All taxa | ENV | 0.261 | 0.1858 | 3.473 | 0.001 | *** |
|  | AEM | 0.1008 | 0.0723 | 3.533 | 0.001 | *** |
|  | EUC | 0.2941 | 0.2089 | 3.453 | 0.001 | *** |
|  | LAND | 0.1402 | 0.0986 | 3.371 | 0.001 | *** |
| Strong fliers | ENV | 0.184 | 0.1304 | 3.438 | 0.001 | *** |
|  | AEM |  |  |  |  | -- |
|  | EUC | 0.0817 | 0.0526 | 2.803 | 0.002 | ** |
|  | LAND | 0.0691 | 0.0395 | 2.338 | 0.020 | * |
| Weak fliers | ENV | 0.2044 | 0.1522 | 3.918 | 0.001 | *** |
|  | AEM | 0.0779 | 0.0635 | 5.407 | 0.001 | *** |
|  | EUC | 0.2635 | 0.2021 | 4.292 | 0.001 | *** |
|  | LAND | 0.1622 | 0.1217 | 4.002 | 0.001 | *** |
| Passive aerial | ENV | 0.1151 | 0.1013 | 8.325 | 0.001 | *** |
|  | AEM | 0.2221 | 0.1845 | 5.902 | 0.001 | *** |
|  | EUC | 0.3722 | 0.3199 | 7.116 | 0.001 | *** |
|  | LAND | 0.1685 | 0.1421 | 6.381 | 0.001 | *** |
| Obligate aquatics | ENV | 0.1581 | 0.1173 | 3.880 | 0.001 | *** |
|  | AEM | 0.1077 | 0.0645 | 2.495 | 0.001 | *** |
|  | EUC | 0.1060 | 0.0777 | 3.737 | 0.001 | *** |
|  | LAND |  |  |  |  | -- |

R<sup>2</sup>, unadjusted coefficient of determination; R<sup>2</sup><sub>adj</sub>, adjusted coefficient of determination; F, pseudo-F statistic; p, permutational significance (999 permutations). Significance: \*\*\* $p \leq 0.001$ , \*\* $p \leq 0.01$ , \* $p \leq 0.05$ ; a dash indicates that no predictor in that block was retained.

The relative importance of predictor blocks shifted across dispersal groups, revealing a dispersal-ability gradient in the dominant predictor block. EUC was the most important block for all taxa (36.9%), weak fliers (37.5%), and passive aerial dispersers (42.8%). For strong fliers, ENV dominated (58.6%), with EUC (23.6%) and LAND (17.8%) together explaining less than half the variance attributed to ENV. Passive aerial dispersers showed the most even distribution among all four blocks (ENV 13.5%, AEM 24.7%, EUC 42.8%, LAND 19.0%), with network connectivity (AEM) contributing a quarter of the explained variance. Obligate aquatics showed the most balanced three-block partition (ENV 45.2%, EUC 29.9%, AEM 24.9%); LAND contributed no explanatory power for this group.

### Variation partitioning

Variation partitioning revealed that the unique fractions of ENV and EUC were significant across most dispersal groups, while the unique contributions of AEM and LAND were more group-specific. The unique environmental fraction was significant for all taxa (R²adj = 0.063, *p* < 0.001), weak fliers (R²adj = 0.053, *p* = 0.006), strong fliers (R²adj = 0.081, *p* = 0.003), and obligate aquatics (R²adj = 0.048, *p* < 0.001), but not for passive aerial dispersers (R²adj = 0.004, *p* = 0.185). The unique EUC fraction was significant for all taxa (R²adj = 0.054, *p* < 0.001), weak fliers (R²adj = 0.053, *p* = 0.004), strong fliers (R²adj = 0.032, *p* = 0.022), and passive aerial dispersers (R²adj = 0.127, *p* = 0.001), but not for obligate aquatics (R²adj = 0.018, *p* = 0.052). The unique AEM fraction was significant only for passive aerial dispersers (R²adj = 0.125, *p* < 0.001) and obligate aquatics (R²adj = 0.036, *p* = 0.008). The unique LAND fraction was significant for all taxa (R²adj = 0.020, *p* = 0.032), weak fliers (R²adj = 0.026, *p* = 0.029), and passive aerial dispersers (R²adj = 0.028, *p* = 0.045), but not for strong fliers (R²adj = 0.013, *p* = 0.124).

### Distance decay patterns

Distance decay relationships (Table 4) were significant but weak across most dispersal groups and distance types, showing that geographic and environmental distance gradients structure community dissimilarity in the Nzoia catchment. EUC produced significant decay relationships for all taxa (Mantel r = 0.112, *p* = 0.003), strong fliers (r = 0.068, *p* = 0.040), passive aerial dispersers (r = 0.143, *p* = 0.005) and obligate aquatics (r = 0.176, *p* = 0.001), but not for weak fliers (r = 0.055, *p* = 0.096). AEM generated significant decay only for passive aerial dispersers (r = 0.165, *p* = 0.005) and obligate aquatics (r = 0.263, *p* < 0.001), the two groups for which AEM explained significant variance, and the strongest network distance decay signal recorded in the dataset (R² = 0.069) was for obligate aquatics. LAND produced significant decay relationships for all taxa (r = 0.130, *p* = 0.017), strong fliers (r = 0.127, *p* = 0.011), passive aerial dispersers (r = 0.097, *p* = 0.049) and obligate aquatics (r = 0.172, *p* < 0.001), but not for weak fliers (r = 0.084, *p* = 0.073). ENV was a significant predictor of dissimilarity only for weak fliers (r = 0.131, *p* = 0.049) and obligate aquatics (r = 0.186, *p* = 0.008), with no significant relationshi*p* detected for strong fliers (r = 0.003, *p* = 0.510) or passive aerial dispersers (r = 0.081, *p* = 0.153).

**Table 4.** Mantel tests of distance decay in macroinvertebrate community dissimilarity against environmental and spatial distance. Tests were run separately for the full assemblage and the four dispersal groups. Community dissimilarity is pairwise Bray-Curtis distance.

| Group | Predictor Block | Mantel r | p-value | Significance |
| --- | --- | --- | --- | --- |
| All taxa | ENV | 0.1525 | 0.036 | * |
|  | AEM | 0.0776 | 0.084 | ns |
|  | EUC | 0.1124 | 0.003 | ** |
|  | LAND | 0.1303 | 0.017 | * |
| Strong fliers | ENV | -0.0032 | 0.510 | ns |
|  | AEM | -0.0168 | 0.607 | ns |
|  | EUC | 0.0679 | 0.040 | * |
|  | LAND | 0.1272 | 0.011 | * |
| Weak fliers | ENV | 0.1312 | 0.049 | * |
|  | AEM | 0.0287 | 0.300 | ns |
|  | EUC | 0.0546 | 0.096 | ns |
|  | LAND | 0.0839 | 0.073 | ns |
| Passive aerial | ENV | 0.0811 | 0.153 | ns |
|  | AEM | 0.1649 | 0.005 | ** |
|  | EUC | 0.1429 | 0.005 | ** |
|  | LAND | 0.0971 | 0.049 | * |
| Obligate aquatics | ENV | 0.1859 | 0.008 | ** |
|  | AEM | 0.2626 | 0.001 | *** |
|  | EUC | 0.1764 | 0.001 | *** |
|  | LAND | 0.1720 | 0.001 | *** |

ENV, environmental distance; EUC, Euclidean distance; LAND, land-use resistance distance; AEM, river-network distance. Mantel $r$ , standardised Mantel correlation (999 permutations). Significance: \*\*\* $p \leq 0.001$ , \*\* $p \leq 0.01$ , \* $p \leq 0.05$ , ns $p > 0.05$ .

### Beta diversity decomposition

Beta diversity decomposition revealed that turnover (beta.SIM) dominated total compositional dissimilarity across most groups, accounting for 94.9 to 96.1% of total beta in all taxa combined, strong fliers, weak fliers, and obligate aquatics (Figure 5). Passive aerial dispersers were the exception, exhibiting the lowest turnover proportion (88.8%) and the highest nestedness component (beta.SNE = 0.101) of any group. In the pairwise distributions, this was the only group in which mean nestedness exceeded mean turnover (Figure 5b). Turnover and nestedness distributions also varied among groups (Figure 5a).

**Figure 5.**
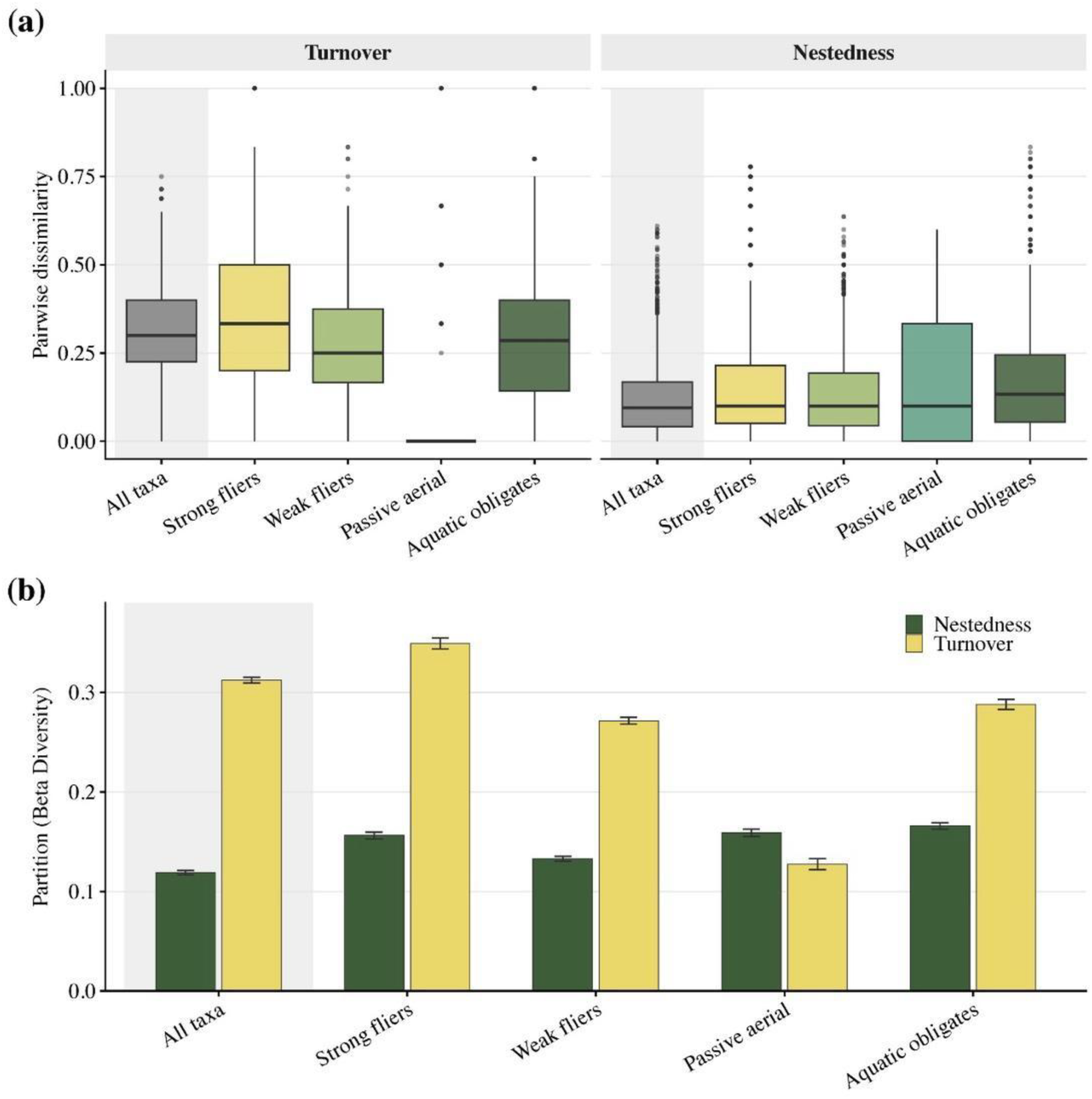
Beta-diversity decomposition of macroinvertebrate community dissimilarity into turnover and nestedness components, shown for the full assemblage and the four dispersal groups. Total Sorensen dissimilarity was partitioned into turnover (Simpson dissimilarity, beta.SIM) and nestedness (beta.SNE). (a) pairwise dissimilarity distributions for each component; (b) mean pairwise turnover and nestedness per group, with error bars showing the standard error of the pairwise distribution.

## Discussion

This study shows that dispersal ability changes how stream macroinvertebrate metacommunities respond to environmental filtering and spatial connectivity. Its main contribution is not that space matters, but that the relevant spatial pathway differs among dispersal groups. The overland, land-use mediated and along-watercourse dispersal pathways were not interchangeable. Each represents a different route by which organisms reach sites in a dendritic landscape.

## Dispersal ability governs the balance between environmental filtering and dispersal

Environmental filtering shaped community composition across all aerial dispersal groups, but its relative importance declined as dispersal ability decreased, while the overland and along-watercourse dispersal pathways became progressively more important. This gradient reflects a central paradigm of metacommunity theory: when organisms disperse freely, local environmental conditions determine species occurrence at each site, so communities reflect habitat quality rather than spatial position (Leibold et al., 2004; Heino et al., 2015; Fuß et al. 2026). Strong fliers were structured primarily by environmental filtering. Their greater aerial mobility likely reduces dispersal limitation, enabling them to reach suitable sites across the catchment, so local physicochemical and habitat conditions then determine establishment and persistence. This pattern is consistent with species sorting and with evidence that environmental gradients explain more variation than spatial separation among strong-dispersing stream macroinvertebrates (Leibold et al., 2004; Sarremejane et al., 2017; Li et al., 2021). Community dissimilarity in this group nevertheless increased with the geographic distance between sites, so the overland dispersal pathway retained a detectable influence on composition even where environmental filtering dominated.

Weak fliers responded in equal measure to environmental filtering and to the overland dispersal pathway, and the land-use mediated pathway added a further independent fraction, consistent with metacommunity theory in which the balance of environmental and spatial control depends on dispersal ability (Leibold et al., 2004; Cañedo-Argüelles et al., 2015). Local environmental conditions selected taxa on the basis of water quality and instream habitat, while overland distance independently limited community similarity, so sites further apart shared fewer taxa irrespective of environmental similarity (Nekola & White, 1999). This pattern is consistent with dispersal limitation, because overland distance constrains weak dispersers more strongly than strong dispersers, and many weak-flying Ephemeroptera, Plecoptera and Trichoptera adults remain close to their natal streams (Petersen et al., 2004; Sarremejane et al., 2017; Li et al., 2021). Community dissimilarity in this group also increased with environmental distance, one of only two groups in which it did so, which supports the independent role of local physicochemical conditions. The independent contribution of the land use mediated pathway shows overland distance alone represented connectivity incompletely, since equal distances differed in permeability to dispersing adults. The composition of the intervening landscape modified the cost of overland movement, likely through differences in vegetation cover, microclimate and the continuity of suitable dispersal corridors (Morán-Ordóñez et al., 2015; Firmiano et al., 2021). The overland contribution to this group was detected by the spatial factors but not by the pairwise distance-decay analysis, so weak-flier assemblages were spatially structured without showing a detectable continuous decline in similarity with distance. Eigenvector-based spatial factors detect spatial structure a pairwise correlation does not (Legendre et al., 2015), and the two analyses are therefore complementary rather than contradictory. Turnover dominated beta diversity in this group, indicating communities remained compositionally distinct among sites, as expected where overland dispersal limitation restricts the exchange of taxa (Chaput-Bardy et al., 2008; 2012; Firmiano et al., 2021; Turunen & Snåre, 2025).

Among dispersal groups, passive aerial dispersers were most strongly structured by spatial connectivity. The overland and along-watercourse pathways each explained a unique component of community composition, whereas environmental conditions explained none. Community dissimilarity in this group increased with both the distance between sites and the distance along the river network, yet showed no increase with environmental distance, consistent with dispersal rather than local physicochemical conditions governing assembly. Two non-exclusive mechanisms account for the downstream signal. First, taxa with weak individual flight capacity are transported by wind along the river corridor, where valley topography channels airflow (Winterbourn et al., 2007; Sukhodolov et al., 2023). Second, larvae drift passively downstream during the aquatic stage, between environmental conditions so movement along the network proceeds independently of adult flight (Bilton et al., 2001; Heino, 2013; Liu et al., 2013). Both mechanisms displace individuals downstream along the same pathway, but we don’t have data to determine which of them dominated, so both remain plausible contributors here. The variance shared and the two pathways arises because water temperature, the only environmental predictor retained for this group, follows the same altitudinal and longitudinal gradient as the spatial variables (Wanderi et al., 2022). Passive aerial dispersers were also the only group in which nestedness exceeded turnover, meaning sites with fewer taxa hosted subsets of the taxa found at richer sites. Continuous delivery of individuals along the watercourse offers one explanation for this pattern, sustaining populations at sites where local physicochemical conditions alone would not predict their occurrence (Li et al., 2021; Liu et al., 2024). Morán-Ordóñez et al. (2015) reported comparable patterns for passive aerial dispersers in Australian drainage basins.

## Land-use resistance and directional river network connectivity impose group-specific spatial constraints

Land-use resistance determined successful dispersal for all aerial groups, but its strength differed with dispersal capacity and was absent entirely for obligate aquatics that disperse exclusively through the river network. For weak fliers and passive aerial dispersers, the land-use matrix actively removed individuals during transit, with cropland and urban areas reducing connectivity of corridors and imposing physiological costs on dispersing individuals (Bilton et al., 2001; Petersen et al., 2004; Carlson et al., 2016; Richmond et al.,2024). CIRCUITSCAPE quantified this cumulative filtering by integrating dispersal cost across all possible terrestrial pathways simultaneously, capturing total landscape resistance rather than resistance along a single assumed route (McRae et al., 2008).

For obligate aquatics, river network connectivity determined which sites were reachable, setting the spatial boundaries of dispersal, while flow directionality shaped the asymmetry of movement within those boundaries. The along-watercourse pathway explained the largest unique fraction of composition in this group, and community dissimilarity increased more steeply with network distance than in any other dispersal group, identifying position within the network as the dominant spatial control on assembly in the Nzoia catchment. Drift supplies the mechanism. Taxa susceptible to water current enter the water column through behavioural or catastrophic displacement and move downstream, so the assemblage at any site reflects local instream conditions together with cumulative input from connected upstream reaches. This directional dependence means degradation at an upstream site propagates downstream through the same connectivity structuring the community. Macroconsumers capable of moving against the current, such as freshwater shrimps and crabs, form an exception, with upstream movement documented in these taxa (Gratwicke, 2004; Covich et al., 2009). Hou et al. (2022) reported comparable directional structuring for aquatically dispersing groups in subtropical Chinese rivers, indicating river network connectivity and flow directionality operate as general determinants of obligate aquatic assembly. Large residual variance across all dispersal groups indicates the measured predictors capture only part of the assembly process. Unmeasured factors, including substrate type, reach-scale hydraulic diversity, riparian canopy cover, biotic interactions, and seasonal flow variability, plausibly contribute, and the residual magnitude matches values reported in other stream macroinvertebrate metacommunity studies (Li et al., 2021; Liu et al., 2023). Family-level taxonomic resolution, necessitated by limited species-level trait databases for Afrotropical invertebrates (Odume et al., 2023), also attenuated trait-environment relationships detectable at finer resolution.

## Group-specific assembly processes require trait-informed management strategies

Understanding how dispersal ability determines community assembly has direct implications for how conservation and management actions should be prioritised in river catchments. Because different dispersal groups are governed by different dispersal pathways, a single management strategy will not be equally effective across the entire macroinvertebrate community. For strong fliers, local physicochemical conditions determined species occurrence and persistence at a site, so improving water and habitat quality is the most appropriate management option for this group. For weak fliers and passive aerial dispersers, the land-use mediated dispersal pathway determines which individuals disperse successfully, adding a constraint that instream restoration alone cannot address. Restoring riparian dispersal corridors and improving the permeability of the land-use matrix between stream sites (Supporting Information Figure S2a) would reduce dispersal attrition during transit and allow these groups to reach restored habitats more effectively (Carlson et al., 2016; Firmiano et al., 2021). For most obligate aquatics, the along-watercourse dispersal pathway governed movement between sites, and the community at any site carries the cumulative biological signal of connected upstream reaches, so maintaining and restoring longitudinal connectivity within the river network is the priority for this group (Lowe et al., 2018; Munzhelele et al., 2024). A network of conservation sites spanning the full altitudinal or longitudinal and land-use gradient will conserve more regional macroinvertebrate diversity than effort concentrated at a single location (Heino et al., 2015). These findings support a growing body of evidence showing that dispersal traits must be incorporated alongside water and habitat quality in freshwater conservation planning for human-modified tropical catchments (Firmiano et al., 2021; Li et al., 2021). Bioassessment frameworks in Afrotropical rivers would benefit from group-specific approaches that account for the dispersal pathway governing each group rather than treating the macroinvertebrate assemblage as a single entity.

## Conclusion

This study shows that dispersal traits determine the balance between environmental filtering and spatial connectivity in Afrotropical stream metacommunities. No single dispersal pathway captured assembly across all groups: the overland, land-use mediated and along-watercourse dispersal pathways each explained different components of community structure. Trait-based connectivity models are therefore necessary for predicting biodiversity responses in human-modified tropical catchments. Future work should test this framework across additional Afrotropical basins and with species-level trait data as regional trait databases improve.

## Acknowledgements

The authors thank Lubanga Lunaligo and Joshua Kimeli of the University of Eldoret for their assistance with field sampling and laboratory analysis. This study was funded by the National Research Fund of Kenya through the KISS Project (FY 2017/2018). Additional support was provided by the European Union through the Intra-Africa Academic Mobility Project, CareForAfrica, which supported the corresponding author’s participation in the study.

## Author Contributions

B.C.C. and F.O.M. conceived the ideas and designed the study. B.C.C., F.O.M., A.S., M.Y. and E.W.W., participated in field sampling. B.C.C., A.S., M.Y., and E.W. processed the samples and performed the taxonomic identifications. B.C.C. curated the dataset and conducted the spatial and statistical analyses. F.O.M. and T.F. provided guidance on the analytical approach and interpretation of the results. B.C.C. prepared the initial draft of the manuscript. All authors critically reviewed and revised the manuscript, approved the final version, and accept responsibility for their respective contributions and the integrity of the work.

## Data Availability Statement

The macroinvertebrate community matrices, environmental and spatial predictor data, and the R code used in the analyses will be archived in the Zenodo Digital Repository on acceptance (DOI: https://zenodo.org/records/21633377).

